# Genetic ablation of *SOD1^G37R^* selectively from corticofugal projection neurons protects corticospinal neurons from degeneration without affecting ALS onset and progression

**DOI:** 10.1101/2020.01.09.900944

**Authors:** Jelena Scekic-Zahirovic, Mathieu Fischer, Geoffrey Stuart Lopez, Thibaut Burg, Marie-Christine Birling, Pascal Kessler, Caroline Rouaux

**Author notes:** Correspondence should be addressed to C.R. Equal contribution (Mathieu Fischer, Geoffrey Stuart Lopez).

## Abstract

While clinical evidence of combined degeneration of the bulbar and spinal motor neurons (MN) together with the corticospinal neurons (CSN) is required to diagnose Amyotrophic Lateral Sclerosis (ALS), preclinical studies have mostly concentrated on MN, leaving aside the CSN and their contribution to ALS onset and progression. Recent studies carried on ALS patients suggest that the disease may initiate in the motor cortex and spread to its projection targets, along the corticofugal axonal projections (including CSN), either via altered neuronal excitability and subsequent excitotoxicity, or via prion-like propagation of misfolded proteins. We recently provided first experimental arguments in favour of the corticofugal hypothesis of ALS, demonstrating that CSN and other subcerebral projection neurons were toxic in a context of ALS. Here, we aimed to determine how CSN may be detrimental to their downstream targets, and what governs their degeneration. To answer these questions, we took advantage of the *FloxedSOD1^G37R^* mouse model of ALS that allows genetic ablation of the mutant transgene in selected cells upon Cre-mediated recombination, and crossed it to the *CrymCreER^T2^* mouse line that we purposely designed to genetically target CSN and other corticofugal projection neurons (CFuPN) populations. We demonstrate that excision of the mutant *SOD1^G37R^* transgene from the CSN is sufficient to prevent their death, suggesting that CSN degeneration mostly relies on cell-intrinsic mechanisms. However, genetic ablation of *SOD1^G37R^* transgene from the corticofugal neurons had no effect on disease onset and survival. The data thus indicate that the toxicity of CFuPN in the context of ALS, and corticofugal propagation of the disease, are not mediated by the presence of misfolded mutant proteins, but more likely by other aspects of the cortical pathology, possibly hyperexcitability.

## Introduction

Combined degeneration of corticospinal neurons (CSN) in the motor cortex, and motor neurons (MN) in the brainstem and spinal cord clinically defines amyotrophic lateral sclerosis (ALS), a terminal disease and the third most frequent neurodegenerative disease after Alzheimer’s and Parkinson’s diseases (Hardiman et al., 2017). If ALS is mostly sporadic, about 10% of patients have a familial history of the disease. The most common causative genes are *C9ORF72*, *SOD1* (the first gene ever linked to ALS), *FUS* and *TARDBP* (Brown and Al-Chalabi, 2017; Chia et al., 2017). These have greatly contributed to the development of cellular and animal models of the disease. Indeed, preclinical studies based on the numerous rodent models that followed the discovery of *SOD1* in particular (Lutz, 2018), and the more recent emergence of induced MN from ALS patients’ iPSC (Guo et al., 2017) have allowed to establish a clear picture of the mechanisms involved in MN degeneration. Contrastingly, little is known at the moment about the mechanisms that govern CSN dysfunction and degeneration, and their role in disease onset and progression. The question is altogether central to ALS research, given the severity of the disease in comparison with diseases that affect only the MN (Kennedy’s disease, adult-onset spinal and muscular atrophy) (Sorenson, 2012), and timely, in the context of a renewed interest for the motor cortex in ALS clinical and preclinical research (Braak et al., 2013; Eisen et al., 2017; Geevasinga et al., 2016).

The pathological hallmark of the great majority of sporadic and familial ALS cases is the presence of intracytoplasmic inclusions of TDP-43, encoded by *TARDBP*, with the exception of familial *SOD1* and *FUS* patients that display respectively SOD1 and FUS aggregates (Farrawell et al., 2015). Detailed examination of the cerebral pathology indicated a greater frequency of aggregates in the motor cortex and its direct targets (Brettschneider et al., 2013), which led to the emergence of the so-called « corticofugal hypothesis » that proposes an origin of the pathology in the motor cortex and a prion-like propagation of abnormal proteins along the corticofugal axonal projections (Braak et al., 2013). Another argument in favour of a cortical origin of the disease is the early cortical hyperexcitability that appears as an intrinsic feature of sporadic and familial amyotrophic lateral sclerosis (ALS) phenotypes, precedes the onset of MN dysfunction and correlates with ensuing MN dysfunction and degeneration (Geevasinga et al., 2016). Interestingly, a recent study demonstrated that neuronal hyperexcitability was sufficient to induce cytoplasmic accumulation of TDP-43 (Weskamp et al., 2019), linking for the first time hyperexcitability and pathology, and further supporting the earliness of cortical hyperexcitability in the cascade of pathogenic events that characterize ALS.

In order to test the corticofugal hypothesis in a controlled manner, we recently generated a mouse model that ubiquitously expresses a mutant of the murine *Sod1* gene, a condition sufficient to develop ALS-like symptoms and premature death (Marques et al., 2019; Ripps et al., 1995), but entirely lacks CSN and other cortical layer V subcerebral projection neurons (SubCerPN) (Molyneaux et al., 2005), that, together with the cortical layer VI corticothalamic neurons, represents the major corticofugal output (Lodato et al., 2014). We demonstrated that absence of CSN and related SubCerPN delayed disease onset, reduced weight loss and motor impairment, and increased the survival of *Sod1^G86R^* mice, suggesting that SubCerPN and CSN in particular may carry detrimental signals to their downstream targets (Burg et al., 2019). Similar results had been previously obtained in the *SOD1^G93A^* rat model of the disease upon knock-down of the transgene selectively in the neuronal populations of the motor cortex (Thomsen et al., 2014).

What these studies did not address is how CSN may be toxic to their downstream targets, and what governs their degeneration. To answer these questions, we took advantage of the *FloxedSOD1^G37R^* mouse model of ALS that allows genetic ablation of the mutant transgene in selected cells upon Cre-mediated recombination (Boillee et al., 2006), and crossed it to the *CrymCreER^T2^* mouse line that we purposely designed to genetically target CSN and other corticofugal projection neuron (CFuPN) populations. We demonstrate that excision of the mutant *SOD1^G37R^* transgene from the CSN is sufficient to prevent their degeneration, but does not impact disease onset and progression. The data thus suggest that CSN degeneration mostly relies on cell-intrinsic mechanisms, and that the toxicity of the CFuPN to their downstream targets does likely not rely on the presence and propagation of misfolded proteins and aggregates.

## Materials and Methods

### Animals

All animal experiments were performed under the supervision of authorized investigators and approved by the local ethical committee of Strasbourg University (CREMEAS, agreements # 00738.01). Animals were housed in the animal facility of the Faculty of Medicine of Strasbourg, with a regular 12 hours light/dark cycle, under constant conditions (21±1°C; 60% humidity). Standard laboratory food and water were accessible ad libitum throughout all experiments. Knock-in *Crym-CreER^T2^* mice were generated by the Institut Clinique de la Souris (ICS, Illkirch, France) by homologous recombination. The *Crym* locus was engineered to include, downstream of the *Crym* coding sequence, a *T2A-CreER^T2^* cassette. Mice were genotyped by PCR of genomic DNA from tail biopsies using the following three primers: GGTTCTTGCGAACCTCATCACTCGT; CCAGGGATGGCAGTGGAAGACC; GCTATCCAACCACAAACATGAATAGGG. TdTomato reporter mice (B6.Cg-Gt(ROSA)26Sortm14(CAG-tdTomato)Hze/J) were obtained from The Jackson Laboratory and *FloxedSOD1^G37R^* were kindly provided by Dr Don W. Cleveland (Boillee et al., 2006). *Crym-CreER^T2^* males were crossed with TdTomato females to generate *Crym-CreER^T2^*/TdTomato to verify *Cre*-mediated recombination of the reporter gene. *Crym-CreER^T2^* males were crossed with *FloxedSOD1^G37R^* females and the F1 generation provided four genotypes of interest: *WTWT; Crym-CreER^T2^/WT; WT/SOD1^G37R^; Crym-CreER^T2^/SOD1^G37R^*. At post-natal day 15, all animals received a single i.p. injection of 4-Hydroxytamoxifen (H6278, Sigma) dissolved in corn oil (C8267, Sigma) at the dose of 0.25 mg/g body weight. Males were used for survival and behavioural studies and histology. Mice were followed weekly, and disease progression was rated according to a clinical scale going from score 4 to 0, as previously described (Rouaux et al., 2007). End-stage animals were euthanized upon reaching score 0, *i.e.* when they were no longer unable to roll over within 10 s after being gently placed on their back (Rouaux et al., 2007). Disease onset was calculated as the time of peak of body weight.

### Semi-quantitative PCR analyses and gel electrophoresis

Cortical layers were microdissected under a SMZ18 microscope (Nikon) from three adjacent 1 mm-thick coronal brain sections obtained using a stainless steel coronal brain matrix (Harvard Apparatus, MA). Cortical layers, spinal cord and brain stem were harvested, rapidly frozen in liquid nitrogen and stored at −80°C until use. Genomic DNA was extracted using the conventional phenol-chloroform/proteinase-K method. 0.5 µg of DNA was amplified under the same conditions used for genotyping (Boillee et al., 2006). The PCR products were fractionated on a 3.0% agarose gel using 1X TAE buffer containing ethidium bromide and were visualized under UV light, and the gels were photographed using a UV gel documentation system. A 1 kbp DNA ladder molecular weight marker (Life Technologies, Rockville, MD) was run on every gel to confirm expected molecular weight of the amplification product. For quantification, band intensities were measured using ImageJ (NIH).

### qPCR analyses

Cortical layers, spinal cord and muscle tibialis anterior were harvested and rapidly frozen in liquid nitrogen and stored at −80°C until analysis. Total RNA was extracted using TRIzol reagent (Invitrogen) and stainless-steel bead in a Tissue Lyser (Qiagen). 1 µg of RNA was reverse transcribed using the iScript cDNA synthesis kit (Bio-Rad). Quantitative PCR (qPCR) was performed with the IQ SYBR Green Supermix (Bio-Rad). Gene expression was normalized by calculating a normalization factor using *Gusb*, *Actb* and *Hsp90ab1* as reference genes for the nervous tissues, and *H1h2bc*, *H2AC* and *H2AX* as references genes for the muscular tissues. The following primer were used:

*hSod1*: CTGTACCAGTGCAGGTCCTC; CCAAGTCTCCAACATGCCTCT

*Crym*: CACATCAATGCTGTTGGAGCC; TCCACATACAGCACCGCTTG

*Gusb*: CGAGTATGGAGCAGACGCAA; AGCCTTCTGGTACTCCTCACT

*Actb*: ATGTGGATCAGCAAGCAGGA; AGCTCAGTAACAGTCCGCCT

*Hsp90ab1*: TACTACTCGGCTTTCCCGTCA; CCTGAAAGGCAAAGGTCTCCA

*AChR-γ:* GAGAGCCACCTCGAAGACAC; GACCAACCTCATCTCCCTGA

*H1h2bc:* AACAAGCGCTCGACCATCA; GAATTCGCTACGGAGGCTTACT

*H2AC:* CAACGACGAGGAGCTCAACAAG; GAAGTTTCCGCAGATTCTGTTGC

*H2AX:* TCCTGCCCAACATCCAGG; TCAGTACTCCTGAGAGGCCTGC

### Motor tests

For all motor tests, mice were trained from 24 to 25 weeks of age, and then followed from 26 weeks of age until death, on a daily basis for general health and neurological symptoms, and once a week for body weight, muscle grip strength inverted grid test and CatWalk. Each motor session consisted of three trials and the results represent the mean of these three trials. Muscle strength was measured using a strength grip meter (Bioseb, BIO-GS3). Four limbs hang test that allows measuring the ability of mice to use sustained limb tension to oppose gravitational force was used as previously described (Burg et al., 2019; Carlson et al., 2016; Scekic-Zahirovic et al., 2017). Mouse gait was analysed with the CatWalk XT (Noldus Information Technology) which generates numerous parameters for quantitative and qualitative analysis of individual footprint and gait (Vergouts et al., 2015). Recordings were performed under the conditions previously described (Burg et al., 2019; Scekic-Zahirovic et al., 2017). Each mouse was allowed to cross freely the recording field of the runway with three independent attempts. Criteria for data collection were *i)* crossing the field in less than 10 seconds and *ii)* a walking speed variation of less than 60%. The six following gait and coordination parameters were analysed: Run Average Speed, Hind limbs Stand Index, Hind limbs Max Contact Max Intensity Mean, Hind limbs Max Contact Mean Intensity Mean, Hind limbs print area and Hind limbs stride length.

### Electromyography

All recordings were performed with a standard EMG apparatus (Dantec) on mice anesthetized with a solution of Ketamine (Imalgène 1000®, Merial; 90mg/kg body weight) and Xylazine (Rompun 2%®, Bayer; 16mg/kg body weight) and kept under a heating mat to maintain physiological body temperature (≈31°C). Electrical activity was monitored on gastrocnemius and tibialis anterior muscles of both limbs for at least 2 minutes, as previously described (Rouaux et al., 2007). For each muscle, a score of 0 (innervated) or 1 (denervated) was given, and the total scores of the four muscles were summed.

### Tail spasticity

Tail spasticity of end stage mice was determined and quantified as previously described (Oussini et al., 2017). Briefly, a monopolar needle electrode (Medtronic, 9013R0312, diameter 0.3 mm) was inserted in segmental tail muscles of paralyzed mice to record reflex activity. Muscles spasms were evoked with mechanical stimulation of the tail. For quantification of the response, value of maximal amplitude (in mV) of LLR signal was measured after stimulation, using ImageJ (NIH).

### *In situ* hybridization

The cDNA clone for mu *Crystallin* (Crym) is a kind gift from the Arlotta lab. Riboprobes were generated as previously described (Arlotta et al., 2005). Nonradioactive *in situ* hybridization was performed on 40 μm vibratome coronal brain sections 0.14 mm anterior to Bregma. Selected sections were mounted on superfrost slides and processed using reported methods (Lodato et al., 2011). Bright field 10X tiles images were acquired with an AxioImager.M2 microscope (Zeiss) equipped a high-resolution B/W camera (Hamamatsu), and run by the ZEN 2 software (Zeiss).

### Retrograde labelling of the CSN

CSN were labelled as previously described (Marques et al., 2019). Briefly, animals were deeply anesthetized with an intraperitoneal injection of Ketamine (Imalgène 1000®, Merial; 120 mg/kg body weight) and Xylazine (Rompun 2%®, Bayer; 16 mg/kg body weight) solution and placed on a heating pad. Laminectomy was performed in the C3-C4 cervical or lumbar (L1-L2) region of the spinal cord, and the dura was punctured using a pulled glass capillary lowered to the dorsal funiculus. Five pressure microinjections of 23 nl of Fluorogold (Fluorchrome) were performed on each side of the dorsal funiculus (Drummond Scientific, Nanoject II). Five days after injection, or upon reaching end-stage, mice were euthanized for histological procedures (see below)

### General histological procedures

Animals were deeply anesthetized with an intraperitoneal injection of Ketamine (Imalgène 1000®, Merial; 120 mg/kg body weight) and Xylazine (Rompun 2%®, Bayer; 16 mg/kg body weight), and transcardially perfused with cold 0.01 M PBS, followed by cold 4% PFA in 0.01 M PBS. Brains, spinal cords, gastrocnemius and tibialis were post-fixed in the same fixative solution overnight (nervous tissues) or for 1 hour (muscles) and stored in PBS 0.1M at 4°C until use. Fixed brains and spinal cords were cut in 40µm thick sections on a vibratome (Leica Biosystems, S2000).

Immunohistochemistry was performed on brain and spinal cord sections at room temperature. Sections were first immersed in 3% hydrogen peroxide (H_2_O_2_) to remove the endogenous peroxidase activity, and washed with phosphate buffered saline (PBS), incubated with 5% horse serum and 0.5% triton X-100 in PBS for 30 min and incubated with primary antibody overnight. After rinsing in PBS, sections were incubated with biotinylated secondary antibody for 2 hours, rinsed in PBS and incubated 1 hour with the Vectasatin ABC Kit (Vector Laboratories, PK7200).

Revelation was performed by incubating the sections in 0.075% 3, 3’-diaminobenzidine tetrahydrochloride (Sigma-Aldrich, D5905) and 0.002% H_2_O_2_ in 50mM Tris HCl. For immunofluorescence, brain and spinal cord sections were heated at 80°C in citrate buffer pH6.0 for 30 min and incubated with 5% horse serum and 0.5% triton X-100 in PBS for 1 hour before being incubated with primary antibody overnight at 4°C. After rinsing in PBS, sections were incubated with Alexa-conjugated secondary antibody for 2 hours, rinsed in PBS and water and mounted with DPX mounting solution (SIGMA, 06522-100ML).

The primary antibodies used in this study are the following: goat anti-Td-Tomato (MyBiosource, MBS448092, 1/100), rabbit anti-Td-Tomato (Rockland, 600-401-379, 1/100), mouse anti-CRYM (Abnova AA215-314, 1/100), rabbit anti-CRE (BioLegend 908001, 1/100), rat anti-CTIP2 (Abcam ab18465, 1/100), rabbit anti-TLE4 (Santa Cruz sc-9125, 1/100), mouse anti-SATB2 (Abcam ab21502, 1/50) mouse anti-PV (Sigma P3088, 1/1000), goat anti-TPH2 antibody (Abcam ab121013, 1/500), goat anti-choline acetyltransferase (Merk Millipore AB144P, 1/25), mouse anti-human SOD1 (MM-0070-P, clone B8H10, 1/100), mouse anti-P62 (Abcam ab56416, 1/100), rabbit anti-GFAP (Dako Z0334, 1/100), goat anti-Iba1 (Abcam ab5076, 1/100); biotinylated (1/500) and Alexa-conjugated (1/500) secondary antibodies were purchased from Jackson. Images were captured using an AxioImager.M2 microscope (Zeiss) equipped with a structured illumination system (Apotome, Zeiss) and a high-resolution B/W camera (Hamamatsu), and run by the ZEN 2 software (Zeiss). Experimenters blinded to the genotypes performed the neuronal counts and NMJ assessments.

### Brain histological analyses

Brains sections from Fluorogold injected animals were mounted on slides with DPX mounting solution (SIGMA, 06522-100ML). Fluorogold*-*positive neurons were manually counted, on both hemispheres, within layer V, from M2 medially to S1 laterally. IBA1 and GFAP intensities were quantified using ImageJ software (NIH, MD, USA). Three regions of interest (ROI) were created and used for all images. The three ROI were placed respectively over the cortical layers II/III and V and over the cingulate cortex for background. Background was substracted from intensities of the cortical layers II/III and V ROI. For each animal 2 pictures corresponding to each hemisphere were analysed. Serotonergic neurons were quantified from individual coronal sections located at Bregma - 4.72 mm.

### Spinal cord histological analyses

Lumbar (L2-L3) spinal motor neurons from two sections separated by 320 μm were mounted on slides, dried at 40°C on a hot plate for 3 hrs and stained with 1% toluidine blue solution for 1 min, rinsed with water, dehydrated and mounted. Two images per section (one per ventral horn) were captured and motor neurons (defined as cells with area larger than 350 μm^2^) were quantified using ImageJ (NIH).

hSOD1, GFAP, IBA1 and P62 intensities were quantified using ImageJ software (NIH). For each image, ROI were manually drawn following the natural border between grey and white matter. Intensities were quantified similarly to IBA1 and GFAP in the brain (see above).

### Muscle histological analyses

Tibialis anterior muscles were dissected into bundles and processed for immunofluorescence using a combination of rabbit anti-synaptophysin (Eurogentec, 1/50) and rabbit anti-neurofilament antibodies (Eurogentec, 1/50) followed by an Alexa-conjugated donkey anti-rabbit 488 (Jackson, 1/1 000), and rhodamine conjugated α-bungarotoxin (Sigma-Aldrich, T0195), as previously described (Marques et al., 2019). On average 100 NMJ per animals were examined.

### Statistical Analysis

Data are presented as mean ± standard error of the mean (SEM). Statistical analyses were performed using GraphPad Prism 6 (GraphPad, CA). Student’s t-test was used for comparison between two groups, and 1-way or 2-way analysis of the variance, followed by Tukey’s multiple comparison post hoc test were applied for three or more groups. For survival and disease onset analysis, animals were evaluated using log-rank test (Mantel-Cox). Fischer exact test was used to determine significant different proportion. Results were considered significant when p<0.05.

## Results

### Generation of Crym-CreER^T2^ mice

In order to test the effects of selective removal of an ALS-causing mutant gene from the CSN in mice, we first sought to generate a mouse line to restrict the expression of the Cre recombinase within the CFuPN populations composed of the layer V subcerebral projection neurons (SubCerPN) that include the discrete population of CSN, and the layer VI corticothalamic neurons. Within the cerebral cortex, expression of *Mu-crystallin* (*Crym*) was found restricted to CFuPN (Arlotta et al., 2005). We further confirmed this pattern of expression by conducting *in situ* hybridization analyses at postnatal days (P) P1 and P15, and in the adult mouse brain (Fig. 1a, and data not shown). At P15, *Crym* expression was detected in the cingulate cortex, the layer V of the motor and somatosensory cortex, the piryform cortex, the Olfactory Tubercule, and, at a lower level, in the layer VI of the cerebral cortex where corticothalamic neurons are located (Fig. 1a). *Crym* expression was also detected in the most medial part of the caudate putamen, and the CA1 and CA2 regions of the hippocampus (Fig. 1a). To verify that *Crym* was not expressed in the spinal cord, where lower motor neurons (MN) are located, we ran qPCR on mRNA extracted from the layer V of the cerebral cortex and the lumbar part of the spinal cord of 4 months old wild-type mice (Fig. 1b). We did not detect *Crym* expression in the spinal cord (Fig. 1b). We thus designed a knock-in mouse line to restrict the tamoxifen-inducible Cre recombinase (CreER^T2^) within the *Crym* locus, downstream of the coding sequence, and separated from it by a T2A sequence. This allowed to maintain *Crym* endogenous expression and to obtain a spatial and temporal expression of *CreER^T2^* similar to that of endogenous *Crym* (Fig. 1c). We then crossed the *Crym-CreER^T2^* line to the TdTomato reporter line (Madisen et al., 2009), administered a single dose of tamoxifen to P15 double transgenic mice, and sacrificed the animals at P21 to verify *Crym-CreER^T2^*-mediated recombination of the reporter gene (Fig. 1d). Analyses of sagittal and coronal sections revealed TdTomato expression throughout the whole cortical layer V, but also within layer VI and in the upper layers of the frontal cortex (Fig. 1d). Similarly to endogenous *Crym* expression, TdTomato-positive cells were also detected in the striatum and the CA1 and CA2 regions of the hippocampus. Scattered Td-Tomato-positive cells were also found in the olfactory bulb and cerebellum. Importantly, no Td-Tomato positive cells could be detected in the brain stem and spinal cord where MN are located. Finally, Td-Tomato positive axons were detected as tracts within the internal capsule, cerebral peduncle and dorsal funiculus of the spinal cord, in accordance with Td-Tomato expression by SubCerPN and CSN, and as diffused axons is the parenchyma of the thalamus, in accordance with Td-Tomato expression in cortical layer VI where corticothalamic neurons are located (Fig. 1d). To further verify the identity of Td-Tomato-positive cells present in the cortical layers V and VI, we ran a series of immunolabelling and colocalization analyses using optical sectioning by structured illumination (Fig. 1e). As expected, Td-Tomato expression was restricted to CRYM-expressing neurons and CRE-expressing cells. Td-Tomato also colocalized with CTIP2, a transcription factor expressed at high levels by layer V SubCerPN, and at lower levels by layer VI corticothalamic neurons, and with TLE4 a transcription factor expressed by layer VI corticothalamic neurons (Molyneaux et al., 2007). Importantly, Td-Tomato did not colocalize with SATB2, a transcription factor that selectively labels callosal projection neurons across all cortical layers (Fame et al., 2011; Molyneaux et al., 2007), nor with Parvalbumin-positive (PV) inhibitory GABAergic interneurons (Tremblay et al., 2016) (Fig. 1e). To further test whether the chosen strategy could induce recombination within the disease-relevant CSN, we further retrogradelly labelled this neuronal population in the *Crym-CreER^T2^*;TdTomato double transgenic mice by injecting of Fluorogold (FG) within the cervical portion of the dorsal funiculus. Microscopic analyses in the cortical layer V of the motor cortex revealed colocalization of TdTomato with FG (Fig. 1f). Finally, we verified the absence of recombination within the spinal cord. As expected, Td-Tomato positive corticospinal tract (CST) was detected in the ventral part of the dorsal funiculus, but no Td-Tomato positive cell could be detected within the white or the grey matter (Fig. 1g). Importantly, ChAT-positive MN of the ventral horn were devoid of Td-Tomato expression (Fig. 1g). Together, the data indicate that *Crym-CreE^T2^* expression faithfully recapitulates *Crym* expression and allows the genetic targeting of CFuPN and CSN in particular.

**Figure 1:**
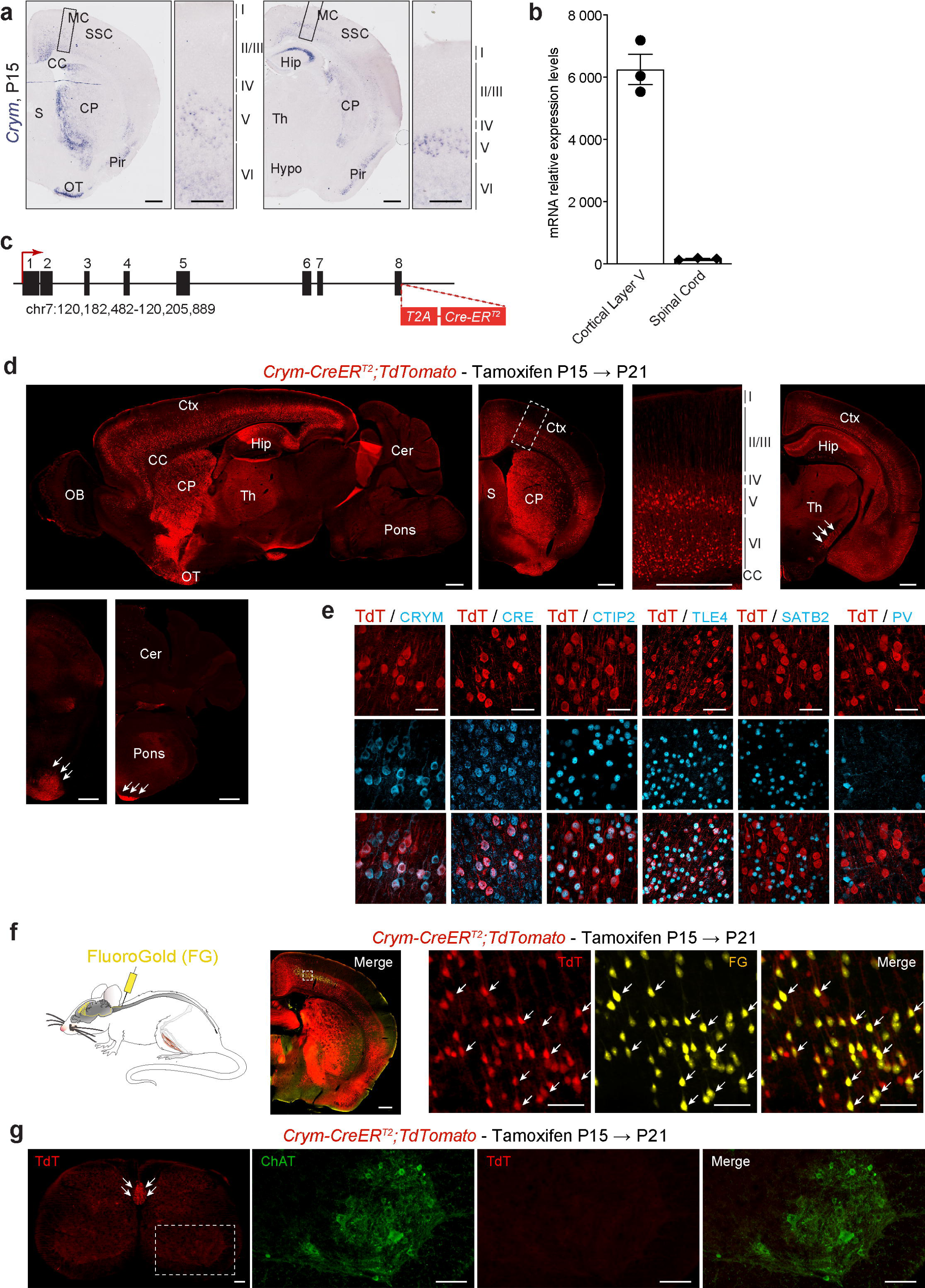
Generation of the knock-in *Crym-CreER^T2^* mouse line. **a** Representative images of *in situ* hybridization on brain coronal sections from a P15 wild-type mouse showing *Crym* expression. Scale bars 500μm and 200µm in close-ups. **b** Graph bar representing RT-qPCR results of relative *Crym* expression in the layer V of the cerebral cortex and in the spinal cord. **c** Schematic representation of the gene targeting strategy to generate the knock-in and tamoxifen-inducible *Crym-CreER^T2^* mouse line. **d** Representative immunofluorescence images of sagittal and coronal sections of brains from P21 *Crym-creER^T2^;TdTomato* double transgenic mice upon induced recombination of the reporter by tamoxifen injection at P15. Arrows indicate Td-Tomato-positive corticospinal tract at different rostro-caudal levels. Scale bar 500μm. **e** Representative optical sectioning images by structured illumination of immunofluorescence showing the colocalization of cortical layer V Td-Tomato-positive cells together with the CRE recombinase, the layer V subcerebal projection neuron markers CRYM and CTIP2, and of cortical layer VI Td-Tomato-positive cells together with the layer VI corticothalamic neuron marker TLE4. Contrastingly no co-localization with the callosal projection neuron marker STAB2 or the interneuron marker PV could be observed. Scale bar 50µm. **f** *Crym-creER^T2^;TdTomato* double transgenic mice received tamoxifen injection at P15 and underwent retrograde labelling of the CSN by injection of FluoroGold into the cervical part of the dorsal funiculus (left schematic). Representative immunofluorescence images of a coronal brain section (right panel) showing Td-Tomato expression and FluoroGold, and their colocalization within the layer V of the motor cortex. Scale bar 500µm and 50µm in the close-ups. **g** Representative immunofluorescence images of a section of the lumbar spinal cord from P21 *Crym-creER^T2^;TdTomato* double transgenic mice showing Td-Tomato-positive CST in the ventral part of the dorsal funiculus, and absence of Td-Tomato expression in the rest of the spinal cord, and more particularly in the ChAT-positive MN of the ventral horn. Scale bar 100µm. MC: motor cortex; SSC: somatosensory cortex; CC: corpus callosum; S: septum; CP:cerebral peduncle; Pir: Piryform cortex; OT: olfactory tubercule; Hip: hippocampus; Th: thalamus; Hypo: hypothalamus; OB:olfactory bulb; Ctx: cortex; Cer: cerebellum.

### Genetic ablation of SOD1^G37R^ from Crym-CreER^T2^ expressing cells

To test the effect of selective removal of an ALS-related transgene from the CSN and other CFuPN, we took advantage of the *Floxed SOD1^G37R^* transgenic mouse line (Boillee et al., 2006) in which the mutant transgene *SOD1^G37R^* can be excised upon Cre-mediated recombination, and crossed it to our newly generated *CrymCre-ER^T2^* line. We obtained 4 genotypes of interest, which we followed over time: *wild-type-non-transgenic* (*WT*), *CrymCreER^T2^-non-transgenic* (*CrymCre*), *wild-type-SOD1^G37R^* (*SOD1*) and *CrymCreER^T2^-SOD1^G37R^* (*CrymSOD1*) (Fig. 2a). All animals received a single dose of Tamoxifen at P15 (Fig. 2a). To verify *SOD1^G37R^* transgene excision in the cortical layer V of *CrymSOD1* animals, compared to their *SOD1* littermates, we microdissected the cortical layers II/III and V, along with the brainstem and spinal cord of four months old, extracted mRNA and genomic DNA and ran respectively RT-qPCR and PCR (Fig. 2b). We observed a significant decrease of *SOD1^G37R^* transgene expression selectively within layer V of the cerebral cortex, and not in the layer II/III or the spinal cord (Fig. 2c). Similarly, we quantified a significant decrease of *SOD1^G37R^* transgene within the genomic DNA extracts from the cortical layer V but not in the cortical layer II/III, the brain stem or the spinal cord (Fig. 2d). As expected, ablation of *SOD1^G37R^* transgene within the cortical layer V is not complete, given the selected expression of the Cre recombinase in only a subpopulation of neurons, *i.e.* the SubCrePN, and not in the other neuronal and glial populations, *i.e.* the excitatory callosal projection neurons, inhibitory interneurons, astrocytes, microglia and oligodendrocytes (Lodato et al., 2014). Together, the data suggest that our strategy to excise *SOD1^G37R^* transgene from CSN, and more broadly SubCerPN and other CFuPN, is efficient.

**Figure 2:**
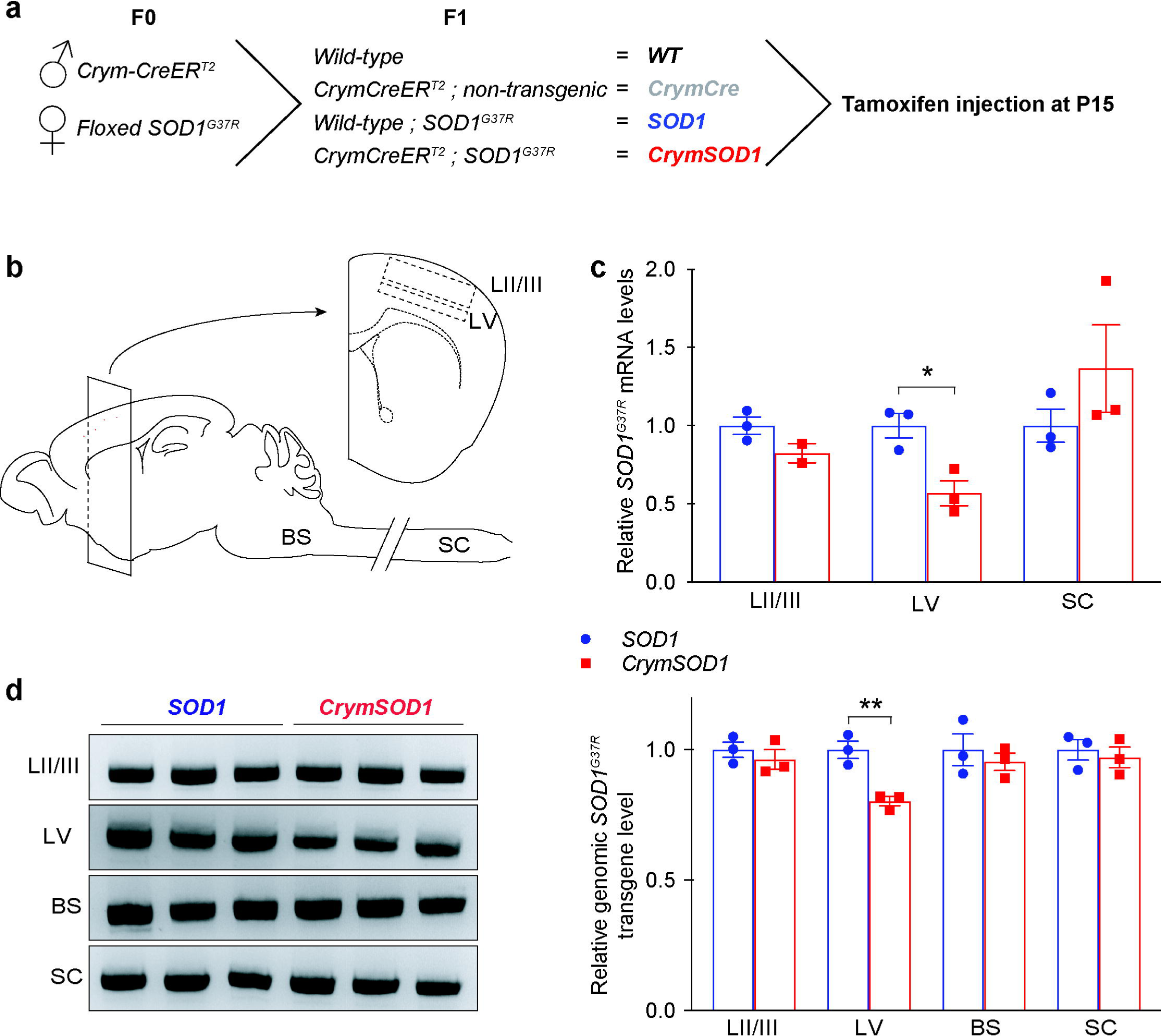
Selective ablation of *SOD1^G37R^* transgene in the cortical layer V. **a** Schematic representation of mice cross-breeding to obtain the four genotypes of interest: *wild-type (WT)*; *CrymCreE^T2^;non-transgenic (CrymCre)*; *wild-type;SOD1^G37R^ (SOD1)*, and *CrymCreE^T2^;SOD1^G37R^ (CrymSOD1)*. **b** Schematic representation of the mouse central nervous system indicating the microdissected areas: layers II/III (LII/III) and V (LV) of the cerebral cortex, brain stem (BS) and the lumbar portion of the spinal cord (SC). **c** Bar graph representing qPCR analyses of the *SOD1^G37R^* transgene expression levels in *SOD1* (blue) and *CrymSOD1* (red) mice. **d** Representative images of agarose gels, and corresponding quantification bar graph, of PCR on genomic DNA to detect the levels of genomic *SOD1^G37R^* transgene. N=3 animals per genotype. Multiple t-test. *p<0.05, **p<0.01.

Selective removal of SOD1^G37R^ from CSN prevents their neurodegeneration without affecting the surrounding reactive gliosis Degeneration of the CSN is a hallmark of ALS and has been reported in different mutant *SOD1* mouse models of the disease (Marques et al., 2019; Ozdinler et al., 2011; Zang and Cheema, 2002). In the *Sod1^G86R^* mouse line, we further demonstrated that lumbar-projecting CSN are more affected than the rest of the population, in accordance with an initial impairment of the hind limbs, and a somatotopic relationship between cortical and spinal neurodegenerations (Marques et al., 2019) To test whether *SOD1^G37R^* excision from the CSN could impact their survival, we retrogradelly labelled lumbar-projecting CSN by injecting the tracer FluoroGold into the lumbar part of the dorsal funiculus of *SOD1* and *CrymSOD1* animals, along with their control littermates *WT* and *CrymCre*. We performed the labelling at two different ages: symptomatic (11 months) and disease end-stage (14-16 months) (Fig. 3a). Microscopic analyses of lumbar projecting CSN revealed a significant loss of this neuronal population in *SOD1* animals compared to controls *WT* and *CrymCre* at a symptomatic stage (24.27 ± 8.16 % loss, p=0.037) and at disease end-stage (21.79 ± 6.60 % loss, p=0.0096) (Fig. 3b,c), indicating that *Floxed-SOD1^G37R^* recapitulated CSN degeneration, like other mutant *SOD1* mouse lines. Excision of *SOD1^G37R^* transgene from CSN and other CFuPN was sufficient to completely prevent the loss of CSN at both ages (*SOD1* vs *CrymSOD1*: p=0.027 at symptomatic stage and p=0.027 at disease end-stage; *WT & CrymCre* vs *CrymSOD1*: p=0.92 at symptomatic stage and p=0.99 at disease end-stage). In order to test whether loss of CSN, or their protection, could impact other cortical pathological hallmarks of the disease, we performed immunolabellings on end-stage *SOD1* and *CrymSOD1* and their aged-matched control littermates to reveal the astrogliosis marker GFAP, and the microgliosis marker IBA1 (Fig. 3d,f). Immunofluorescence analysis revealed an overall mild increase of GFAP and IBA1 immunoreactivity, that appeared as discrete patches, in *SOD1* and *CrymSOD1* compared to control (Fig. 3d,f). Because increased GFAP and IBA1 immunoreactivity seemed more particularly restricted to the cortical layer V, we further quantified their intensities within this layer, along with the cortical layer II/III (Fig. 3e,g). We observed that GFAP immunoreactivity was not significantly different across genotypes within the cortical layer II/III. Contrastingly, *SOD1* and *CrymSOD1* animals presented a significantly higher GFAP immunoreactivity within the cortical layer V, in comparison with their *WT* and *CrymCre* littermates (*SOD1*: increase of 497 ± 53.68 % compared to *WT & CrymCre*, p<0.0001; *CrymSOD1*: increase of 479 ± 53.68 % compared to *WT & CrymCre*, p<0.0001), but no significant difference could be observed between *SOD1* and *CrymSOD1* animals (p=0.9394) (Fig. 3.e). Similarly, no difference in IBA1 immunoreactivity could be observed across genotypes within the cortical layer II/III, while increased of immunoreactivity in cortical layer V of *SOD1* and *CrymSOD1* animals compared to *WT & CrymCre* could be measured (*SOD1*: increase of 199 ± 35.27 % compared to *WT & CrymCre*, p=0.0198; *CrymSOD1*: increase of 213.57 ± 35.27 % compared to *WT & CrymCre*, p=0.0198), without significant difference between *SOD1* and *CrymSOD1* animals (p=0.8738) (Fig. 3.f). Taken in their whole, the data suggest that *SOD1^G37R^* expression by CSN is sufficient to induce their degeneration in a cell-autonomous manner. In addition, CSN degeneration and local (cortical layer V) reactive gliosis appear to be independent from each other: reactive gliosis is neither a cause nor a consequence of CSN degeneration.

**Figure 3:**
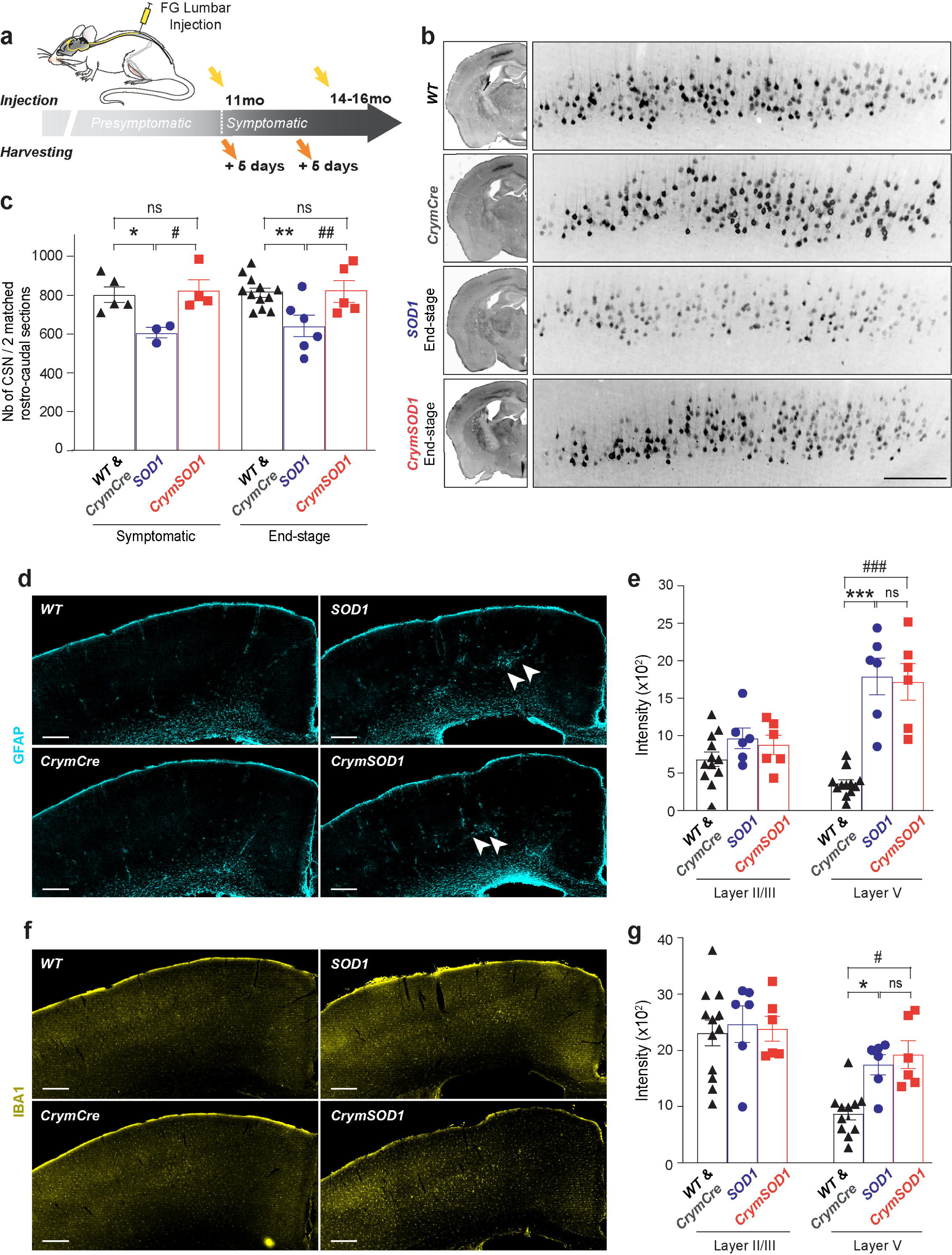
Selective ablation of *SOD1^G37R^* from SubCerPN rescues loss of CSN, but does not prevent reactive gliosis. **a** Schematic of the experimental strategy. **b** Representative negative fluorescence images of the brain hemisphere (left panels) and cerebral cortex (enlarged insets, right panels) of *WT*, *CrymCre*, *SOD1* and *CrymSOD1* mice showing CSN marked upon retrograde labelling with FG at disease end stage. **c** Bar graph presenting the quantification of FG-labelled CSN at symptomatic and disease end stage. N=5 *WT* and *CrymCre*, 3 *SOD1* and 4 *CrymSOD1* at symptomatic stage and N=12 *WT* and *CrymCre*, 6 *SOD1* and 5 *CrymSOD1* at disease end stage. * or ^#^p<0.05 and ** or ^##^p< 0.01 in 1-way ANOVA followed by a Tukey’s multiple comparisons test. **d,e** Representative images of the cerebral cortex *WT*, *CrymCre*, *SOD1* and *CrymSOD1* mice showing GFAP (**d**, blue) and IBA1 (**e**, yellow) immunoreactivity at disease end stage. **e,g** Bar graphs representing the averaged intensity of GFAP (**e**) and IBA1 (**g**) signals in cortical layers II/III and V. N=12 *WT* and *CrymCre*, 6 *SOD1* and 6 *CrymSOD1*. * or ^#^p<0.05, *** or ^###^p<0.001 and ns: non-significant in 2-way ANOVA followed by Tukey’s multiple comparisons test.

### Selective removal of SOD1^G37R^ from the CSN limits spasticity

Degeneration of CSN (or upper motor neurons) leads to the development of the “upper motor neuron syndrome” characterized by decreased motor control, muscular weakness, altered muscle tone, clonus, spasticity and other manifestations of hyperreflexia (Ivanhoe and Reistetter, 2004; Purves et al., 2004; van Es et al., 2017). To test whether maintenance of the CSN population could impact spasticity, we recorded the long lasting tail muscle activity following cutaneous stimulation in awake and fully paralyzed *SOD1* and *CrymSOD1* animals prior to harvesting (Bennett et al., 1999; Dentel et al., 2013; Oussini et al., 2017) (Fig. 4a-c). Within the *CrymSOD1* group, frequency of spastic mice was lower than within the *SOD1* group, but the difference did not reach significance (76.92% vs 90.91% respectively, p=0.5963) (Fig. 4b), however, the amplitude of the response was significantly smaller within the *CrymSOD1* group than within the *SOD1* one (0.3108 ± 0.0492 vs 0.4870 ± 0.0654, p=0.0452), suggesting that *SOD1^G37R^* excision from the CSN and maintenance of the CSN population was sufficient to limit spasticity (Fig. 4c).

**Figure 4:**
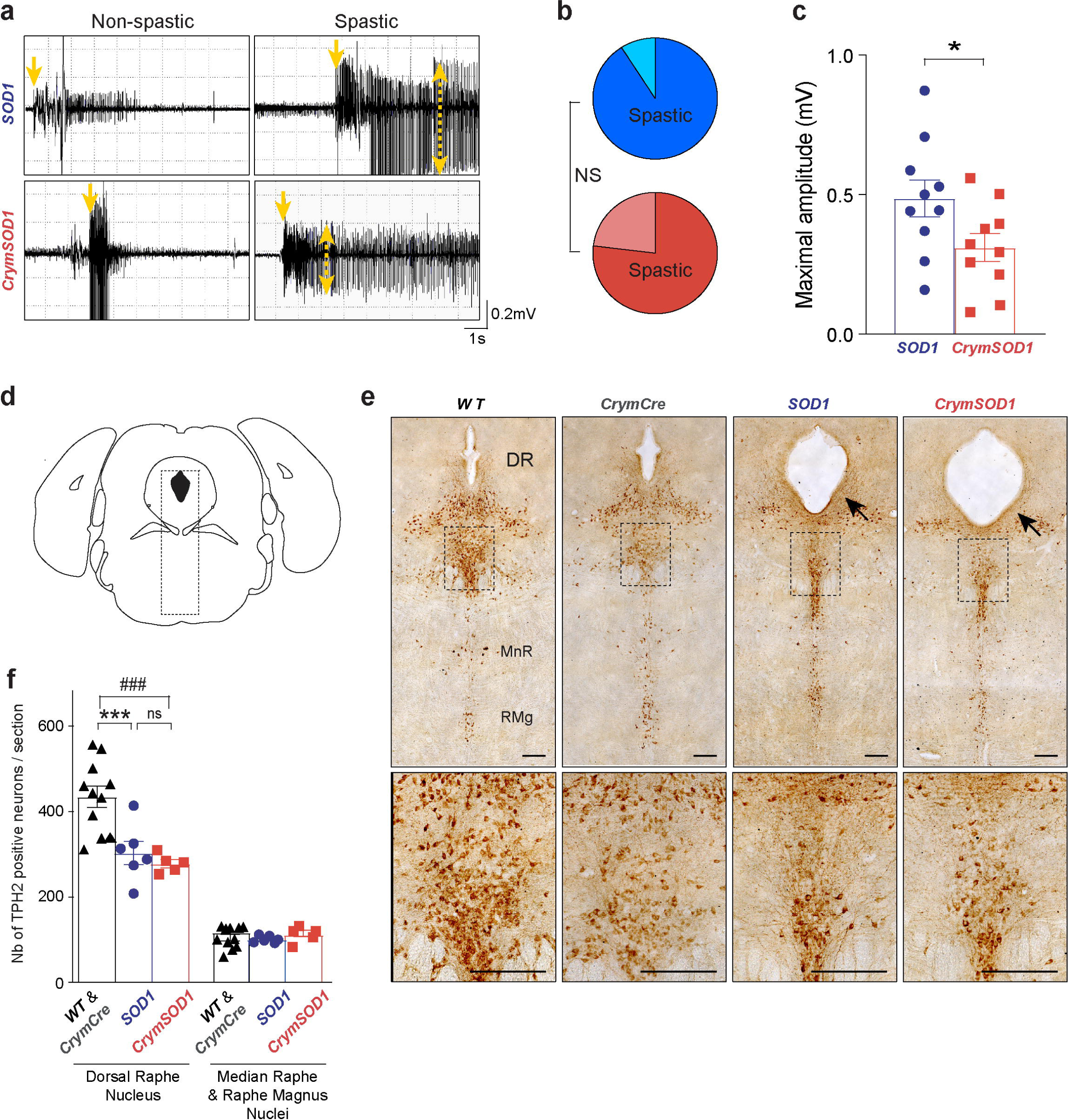
Selective deletion of *SOD1^G37R^* from CSN limits spasticity. **a** Representative EMG recordings of tail’s muscle long lasting reflex (LLR) detected in end stage awake mice *SOD1* (N=11), and *CrymSOD1* (N=13) presenting a complete paralysis of hind limbs. Spasticity-related tail LLR is observed upon stimulation (yellow single head arrows) in subgroups of animals, and amplitude of the response can be recorded (yellow single head arrows). **b** Pie charts representing the frequency of animals with a tail LLR amongst the *SOD1* (light blue: not spastic, dark blue: spastic) and *CrymSOD1* mice (light red: not spastic, dark red: spastic). Difference between the two genotypes is not significant, Fischer’s exact test. **c** Graph bar representing the LLR maximal amplitudes for spastic *SOD1* (N=10) and *CrymSOD1* (N=10) mice. Student unpaired t-test. **d** Schematic of the coronal section selected for TPH2-positive neurons labeling and counting (Bregma –4.84 mm). **e** Representative images of TPH2 immunoreactivity in the brainstem of end stage mice. (DR: dorsal raphe, MnR: median raphe, RMg raphe magnus). Arrowheads indicate the enlargement of the aqueduct (Sylvius) in *SOD1* and *CrymSOD1* animals compared to *WT* and *CrymCre* control animals. Scale bars 200μm. **f** Bar graph representing the average number of TPH2-positive neurons in the dorsal raphe nucleus and the median raphe and raphe magnus nuclei respectively. N= 12 *WT* and *CrymCre*, 6 *SOD1* and 5 *CrymSOD1*. 2-way ANOVA followed by a Tukey’s multiple comparisons test. *p<0.05, *** or ^###^p<0.001 and ns: non-significant.

In the *FloxedSOD1^G37R^* mouse model, spasticity has previously been linked to serotonergic neuron degeneration in the raphe nuclei (Dentel et al., 2013; Oussini et al., 2017). We thus tested whether the decreased levels of spastic response that we recorded with the tail LLR could be at least partly attributed to a limited degeneration of serotonergic neurons. We quantified the number of TPH2-positive neurons present in the dorsal raphe nucleus (DR) and the median raphe and raphe magnus nuclei of end-stage *SOD1* and *CrymSOD1* and their control littermates, *WT* & *CrymCre* (Fig. 4d-f). We did not detect any difference in the median raphe and raphe magnus nuclei across genotypes (Fig. 4e,f). However, we observed a significant loss of TPH2-positive neurons of *SOD1* and *CrymSOD1* animals compared to *WT & CrymCre* animals in the dorsal raphe nucleus (*SOD1*: loss of 30.14 ± 6.02, p<0.0001 and *CrymSOD1*: loss of 35.98 ± 6.40; p<0.0001) (Fig. 4e,f). This loss was accompanied by an enlargement of the aqueduct in the *SOD1* and *CrymSOD1* mice compared to *WT & CrymCre* mice (Fig. 4e, arrows). However, no significant difference in the number of dorsal raphe nucleus TPH2-positive neurons could be observed between *SOD1* and *CrymSOD1* animals (p=0.7057) (Fig. 4f). Together, the data suggest that in the *FloxedSOD1^G37R^* mouse line, intensity of the spastic tail LLR can be at least partly attributed to CSN dysfunction and/or degeneration, and not only to serotonergic neuron degeneration.

### Maintenance of the CSN population does not impact ALS-like symptoms onset and progression

Next, we sought to test the effect of CFuPN-selective ablation of *SOD1^G37R^* on the manifestation of ALS-like symptoms in our mice, *i.e.* weight loss, disease onset, premature death and motor impairment (Fig. 5). As already reported (Boillee et al., 2006), *SOD1* mice stopped gaining weight (Fig. 5a), started progressively losing weight, which we considered to be the onset of the disease from 160 days on (Fig. 5b), and died prematurely from 410 days on (Fig. 5c). While *CrymSOD1* animals were slightly but significantly lighter than their *SOD1* littermates (Fig. 5a, p=0.0121), we did not detect any difference between *SOD1* and *CrymSOD1* animals regarding disease onset (Fig. 5b p=0.4527) and survival (Fig. 5c p=0.2152). Motor performance analyses revealed no difference between the *SOD1* and *CrymSOD1* groups regarding the evolution of the grip strength of the animals (Fig. 5d, p=0.1434), or the time when it started decreasing (Fig. 5e, p=0.1114). However, *CrymSOD1* animals performed significantly better than their *SOD1* littermates on the inverted grid test with a longer latency to fall (Fig. 5f, p<0.0001), a delay of the age at which animals had lost 25% (Fig. 5g, p=0.0004) or 50% of their initial latency to fall (Fig. 5h, p=0.0335), and a higher holding impulse (Fig. 5i, p=0.0009). Contrastingly, Catwalk analyses of the gait revealed no difference between *SOD1* and *CrymSOD1* animals regarding the run average speed (Fig. 5j, p=0.3140), the hind limbs “stand index” (Fig. 5k, p=0.0989), “max contact max intensity mean” (Fig. 5l, p=0.3238), “max contact mean intensity mean” (Fig. 5m, p=0.2483), “print area” (Fig. 5n, p=0.9311) and “stride length” (Fig. 5o, p=0.2803). Taken in their whole, the data indicate that deletion of *SOD1^G37R^* from the CFuPN, and maintenance of the CSN population has no impact on disease onset, and survival, and only ameliorates discrete motor functions.

**Figure 5:**
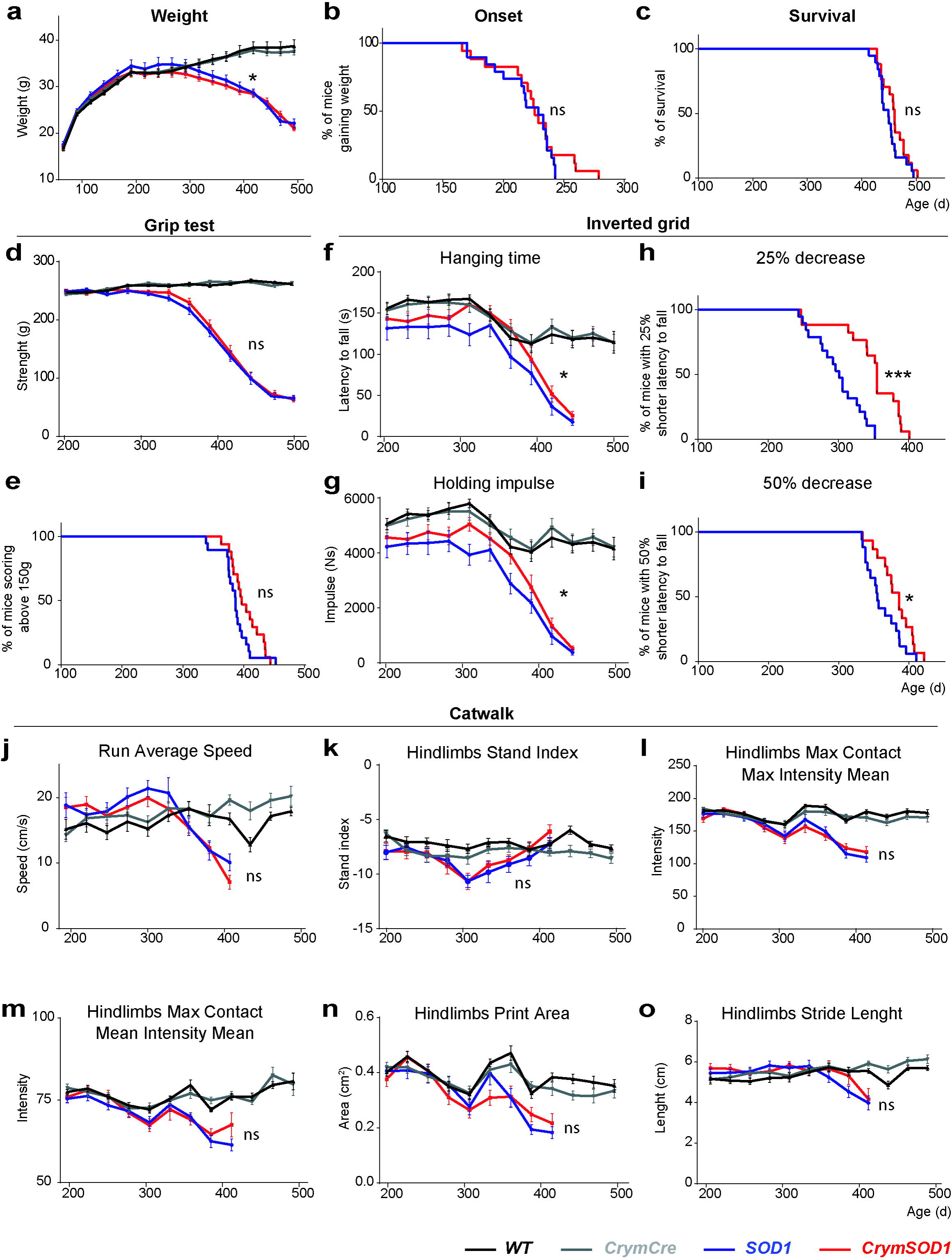
CSN rescue does not impact disease onset or survival. **a** Graphical representation of the longitudinal follow up of the body weight for all genotypes. **b,c** Kaplan-Meier plots of disease onset (age when animals reach their weight peak) and survival (age at death) in days, for *SOD1* (blue) and *CrymSOD1* mice (red). ns: non significant, Log-rank test (Mantel-Cox). **d,e** Graphical representation of the evolution of the grip strength of the animals over time (**d**), and Kaplan-Meyer representation of the percentage of animals presenting a grip strength above 150g (**e**). **f,g,h,i** Graphical representations of the latency to fall from an inverted grid (**f**) and the corresponding holding impulse (**g**) and Kaplan-Meyer representations of the percentage of animals that still present at least 75% (**h**) or 50% of their initial latency (**i**) on that same test. **j,k,l,m,n,o** Graphical representations of selected gait parameters recorded on CatWalk. For all data, *WT* (N=20), *CrymCre* (N=20), *SOD1* (N=15), and *CrymSOD1* (N=14). Data were analyzed by two-way anova (**a,d,f,g,j-o**) or log rank Mantel–Cox (**b,c,e,h,i**) *p<0.05, ***p<0.001, ns: non significant.

### Maintenance of the CSN population does not prevent or delay muscle denervation or spinal pathology

Next, we tested whether ablation of the *SOD1^G31R^* transgene from the CSN and related corticofugal neurons had an impact on the muscular and spinal manifestations of the disease. We first ran electromyographic recordings on the gastrocnemius and tibialis muscles of both hind limbs of end-stage *SOD1* and *CrymSOD1* animals and their *WT* and *CrymCre* age-matched control littermates (Fig. 6a,b). Innervated muscles were given a score of 0, denervated muscles a score of 1 and the sum of the score of the four muscles was calculated for each animal (Fig. 6b). Denervation was clearly observed in *SOD1* and *CrymSOD1* mice compared to the *WT* and *CrymCre* controls, but no significant difference was observed between *SOD1* and *CrymSOD1* mice (*SOD1*: 3.75 ± 0.179; *CrymSOD1*: 3.21 ± 0.317; p=0.0809; Fig. 6b). To further assess the innervation status of the muscles, we labelled the neuromuscular junctions (NMJ) of the tibialis anterior muscle of end-stage *SOD1* and *CrymSOD1* mice and controls *WT* and *CrymCre* (Fig. 6c). Rating of the NMJ integrity into three categories, innervated, partly denervated and fully denervated, confirmed the vast denervation of the *SOD1* and *CrymSOD1* muscles compared to controls, and indicated that *SOD1* and *CrymSOD1* animals presented similar levels of denervation (partially or fully denervated NMJ: *SOD1* 93.15 ± 2.87; *CrymSOD1*: 94.19 ± 2.05; p=0.9474, Fig. 6d). In accordance with this result, qPCR analyses of the tibialis anterior muscle revealed important up-regulation of the expression gamma subunit of the acetylcholine receptor (*AchRγ*), a typical molecular marker of denervation, in end-stage *SOD1* and *CrymSOD1* mice compared to controls (*WT* and *CrymCre*: 1.26 ± 0.64; *SOD1*: 311.98 ± 106.83, p=0.0418; *CrymSOD1*: 439.31 ± 109.37, p=0.046; *SOD1* vs *CrymSOD1* p=0.5730; Fig. 6e). Similarly, quantification of the MN present in the lumbar part of the spinal cord upon toluidine blue labelling indicated an important loss of MN in *SOD1* and *CrymSOD1* animals compared to control (*WT* and *CrymCre*: 9.08 ± 0.71; *SOD1*: 1.70 ± 0.25; *CrymSOD1*: 1.58 ± 0.25) with a similar extent for *SOD1* and *CrymSOD1* mice (p=0.9929) (Fig. 6f,g). Spinal MN degeneration upon *SOD1^G37R^* overexpression in *SOD1* and *CrymSOD1* mice was accompanied with increased levels of hSOD1 protein that appeared as aggregates, together with astrocytosis, microgliosis and altered autophagy as revealed by increased immunolabelling of GFAP IBA1 and P62 respectively (Fig. 7a,c,e,g). Quantification of the fluorescence intensity of these different pathology markers revealed again similar extents of pathology between *SOD1* and *CrymSOD1* animals, both in the white and grey matter (Fig. 7b,d,f,h) with the exception of GFAP that appeared less intense in the white matter of *CymSOD1* mice than in *SOD1* mice (*SOD1*: 19 151.82 ± 804.98; *CrymSOD1*: 16 394.20 ± 1 011.34; p=0.0213). Together, the data indicate that if genetic ablation of *SOD1^G37R^* from the corticofugal neurons is beneficial to CSN survival, it has virtually no impact on ALS pathological hallmarks.

**Figure 6:**
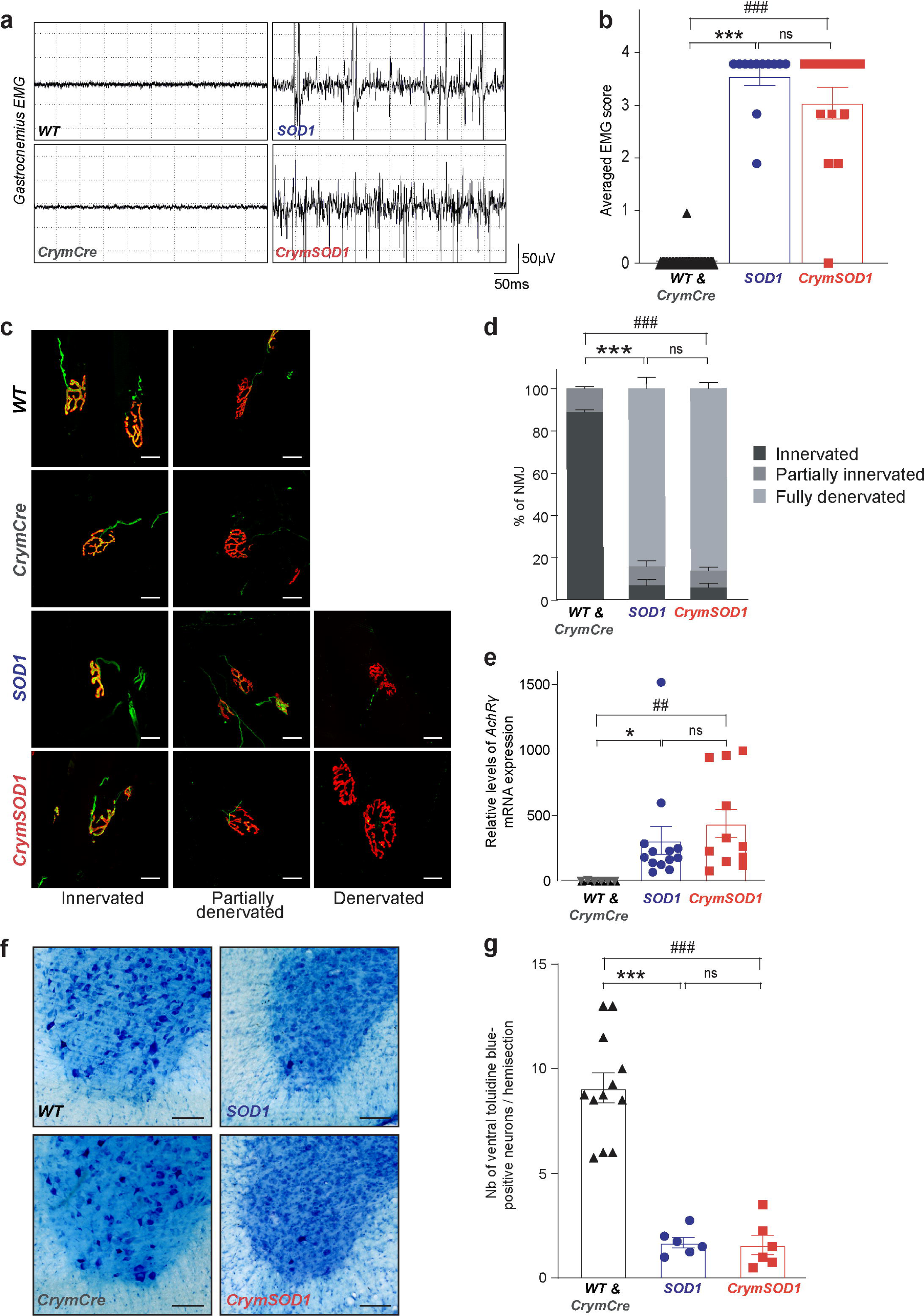
Improved survival of CSN does not prevents denervation nor MN degeneration. **a** Traces of EMG recording of the gastrocnemius muscle of end staged, deeply anaesthetized *SOD1* and *CrymSOD1* animals. **b** Bar graph presenting the EMG score calculated for *SOD1* and *CrymSOD1* mice. N=20 *WT*, 18 *CrymCre*, 12 *SOD1* and 14 *CrymSOD1*. ***p<0.001 and ns: non-significant in 1-way ANOVA followed by a Tukey’s multiple comparisons test. **c** Representative maximum intensity projection images of z-stacks of typically innervated, partially or fully denervated neuromuscular junctions (NMJ) from end-stage *SOD1* and *CrymSOD1* mice and their aged-matched *WT* and *CrymCre* littermates. **d** Bar graph representing the average proportions of innervated (dark grey), partly denervated (medium grey) and fully denervated (light grey) NMJ for each genotype; 2-way ANOVA followed by Tukey multiple comparisons test; N =6 animals per genotype. ***p<0.001 and ns: non-significant. Scale bars 20μm. **e** Bar graph representing qPCR analyses of the *AchRγ* (gamma) mRNA expression levels in *SOD1* (blue) and *CrymSOD1* (red) mice. N=6 per genotype. *p<0.05, ^##^p<0.05 and ns: non-significant in 1-way ANOVA followed by a Tukey’s multiple comparisons test. **f** Representative images of toluidine bleu staining in *WT*, *CrymCre*, *SOD1*, and *CrymSOD1* mice. Scale bars 100μm. **g.** Bar graphs showing mean numbers of spinal motor neuron per hemi-section for all genotypes. N=6 per genotype. *** or ^###^p<0.001 and ns: non-significant in 1-way ANOVA followed by a Tukey’s multiple comparisons test.

**Figure 7:**
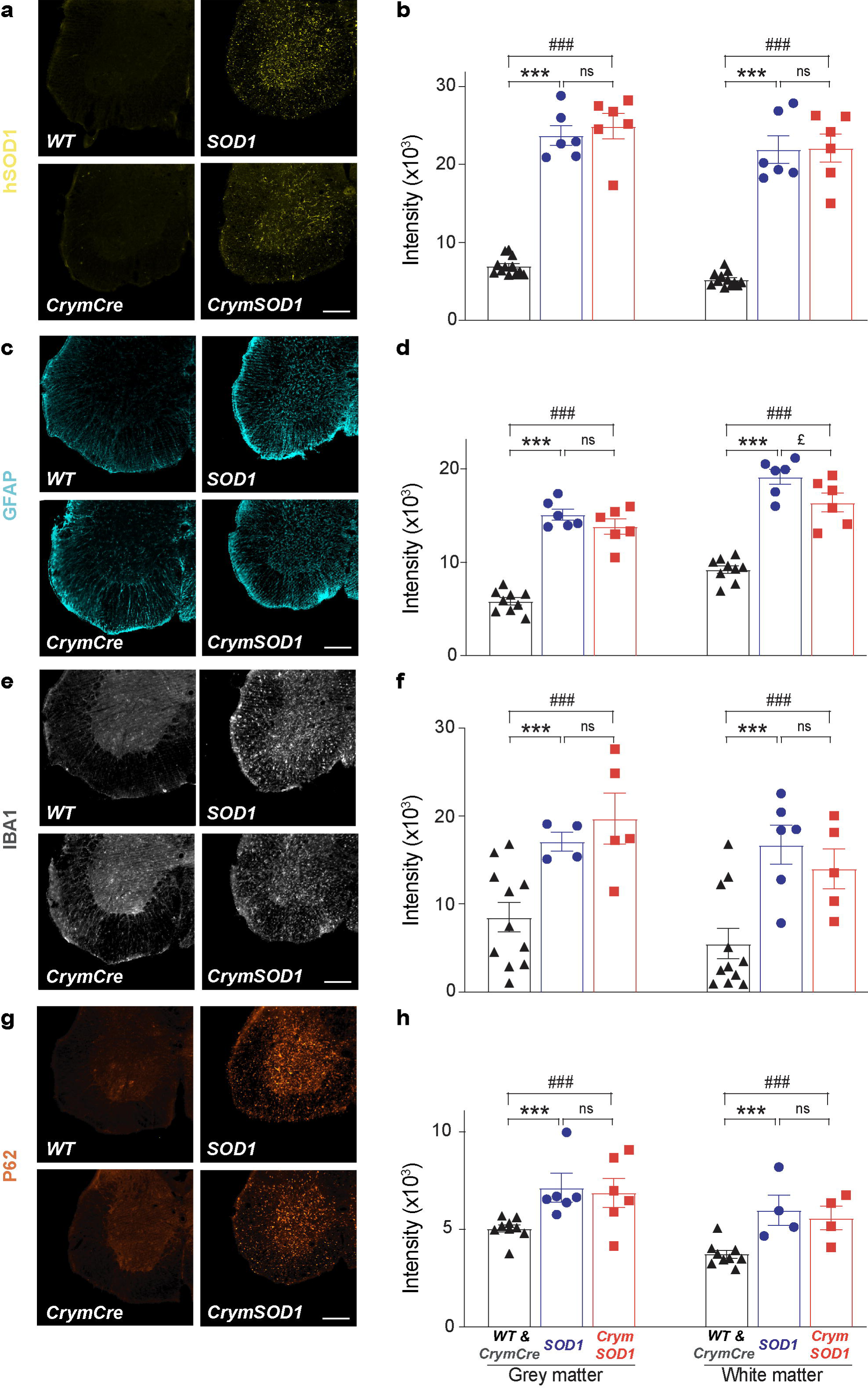
Improved survival of CSN does not modify key pathological hallmarks of ALS in the spinal cord. **a,c,e,g** Representative fluorescence images of spinal cord hemi-sections showing human SOD1 (hSOD1, yellow, **a**), GFAP (cyan, **c**), IBA1 (white, **e**) and P62 (orange, **g**) pathology in end stage *SOD1*, and *CrymSOD1* mice and their age-matched *WT* and *CrymCre* littermates. Scale bar 200µm. **b,d,f,h** Bar graphs representing the average intensity of the hSOD1 (**b**), GFAP (**d**), IBA1 (**f**) and P62 (**h**) specific signal in the grey and white matters of the spinal cord. ^£^p<0.05, *** or ^###^p<0.001 and ns: non-significant in 2-way ANOVA followed by Tukey’s multiple comparisons test.

## Discussion

We previously demonstrated that absence of CSN and other SubCerPN was beneficial in a mouse model of ALS, suggesting that these populations of CFuPN were detrimental in a context of ALS, and bringing first experimental evidence of the corticofugal hypothesis of the disease (Burg et al., 2019). In the present study, we thought to determine how cell-autonomous expression of a mutant toxic transgene *i)* could impact CSN survival, and *ii)* could contribute to the detrimental effect carried by the CFuPN in the peculiar context of ALS. We thus generated a *CrymCreER^T2^* mouse line in order to selectively allow expression of the *Cre* recombinase in CFuPN and excision of the toxic *SOD1^G37R^* transgene (Boillee et al., 2006) from this disease-relevant neuronal population. We demonstrated that genetic ablation of the mutant *SOD1^G37R^* transgene from the CFuPN was sufficient to prevent the loss of CSN, suggesting that CSN degeneration mostly relies on cell-intrinsic mechanisms. However, genetic ablation of *SOD1 ^G37R^* from CFuPN had no effect on disease onset and survival, indicating that toxicity of the CFuPN in ALS, and corticofugal propagation of the disease are probably not mediated by the presence and possible propagation of misfolded mutant proteins, but more likely by other aspects of the cortical pathology, one possibility being cortical hyperexcitability.

### Floxed SOD1^G37R^ mice recapitulate loss of CSN

Progressive degeneration of the CSN translates into a series of symptoms grouped under the term of upper motor neuron syndrome, a hallmark of the clinical manifestation of ALS (Ivanhoe and Reistetter, 2004; Purves et al., 2004). Post mortem analyses of the motor cortex of ALS patients confirms the loss of CSN, or Betz cells (Charcot, 1874; Nihei and Kowall, 1993). Similarly numerous rodent models of the disease recapitulate CSN degeneration such as the *C9-BAC* mice (Liu et al., 2016), the *hPFN1^G118V^* mice (Fil et al., 2016), the *TDP43^A315V^* mice (Herdewyn et al., 2014; Wegorzewska et al., 2009), and the *TDP43^G298S^* mice (Müller et al., 2019) and the *SOD1^G93A^* rats (Thomsen et al., 2014), and *SOD1* mouse models also display progressive loss of CSN that starts pre-symptomatically (Marques et al., 2019; Ozdinler et al., 2011; Yasvoina et al., 2013; Zang and Cheema, 2002). In the *Sod1^G86R^* mouse model of the disease, we further demonstrated that CSN start degenerating long before first signs of MN degeneration can be observed and that CSN and MN degeneration are somatotopically related. Indeed, in the cerebral cortex, lumbar-projecting CSN are affected earlier than the rest of the population, similarly to lumbar MN being affected earlier than the rest of the population in this model (Marques et al., 2019). Here, by sampling the subpopulation of lumbar-projecting CSN, we demonstrated that the *FloxedSOD1^G37R^* mice recapitulate CSN degeneration as well.

### In Floxed SOD1^G37R^ mice, CSN degeneration follows cell-autonomous mechanisms

Genetic ablation of the *SOD1^G37R^* transgene was sufficient to prevent loss of CSN suggesting that CSN degeneration was independent from its *SOD1^G37R^*-expressing environment, *i.e.* other cortical neurons, astrocytes, microglia, oligodendrocytes along their axons and final neuronal targets in the spinal cord, in accordance with strictly cell-autonomous mechanisms. Similar cell-autonomous mechanisms were described for the brainstem serotonergic neurons (Oussini et al., 2017), but not for the spinal MN whose survival rather reflects combination of cell-autonomous and non-cell autonomous mechanism (Boillee et al., 2006). Thus, *SOD1^G37R^* transgene expression differentially affects neuronal survival depending on the nature of the neuronal population. Interestingly, reactive astrogliosis and microgliosis were very moderate in the motor cortex of the *FloxedSOD1^G37R^* animals in comparison with their spinal cord. Cortical astrogliosis appeared as discrete patches, while microgliosis rather corresponded to a two-fold increase of IBA1 expression, and these were selectively found in the layer V of the motor cortex were CSN reside. Surprisingly, maintenance of the CSN population by selective removal of the *SOD1^G37R^* transgene had no impact on reactive gliosis, suggesting that astrogliosis and microgliosis were not secondary to CSN degeneration. Similarly, maintenance of astrogliosis and microgliosis in the cortical layer V of the *CrymSOD1* animals at similar levels than the *SOD1* animals did not prevent the cell-autonomous beneficial effect of *SOD1^G37R^* ablation on CSN survival, suggesting that CSN degeneration is not secondary to reactive gliosis either. Overall, this suggests that CSN degeneration and cortical reactive gliosis are rather independent from each other. The precise co-localization, within the cortical layer V, of CSN degeneration and reactive gliosis remains nevertheless quite remarkable. These results are in opposition with what has been described so far in the spinal cord, *i.e.* the well-described interaction between cell-autonomous and non-cell autonomous mechanisms on spinal MN survival, and the toxic role of astrocytes, microglia and Schwann cells on MN (reviewed in (Ilieva et al., 2009; Lee et al., 2016; Serio and Patani, 2017). The discrepancy between cortical and spinal glial contribution to neighbouring CSN and MN survival may arise from the level of gliosis itself, rather moderate in the cerebral cortex and important in the spinal cord, or from the actual nature of the targeted neuronal populations, which have little in common besides being targeted by ALS. Together, the data strongly suggest that the mechanisms that govern MN degeneration may not be directly transposable to CSN degeneration, and that it is worth further investigating these aspects, as we recently started (Marques et al., 2019).

### Of possible mechanisms of disease propagation by CFuPN

Several clinical features support a cortical origin of ALS (Eisen et al., 2017; Geevasinga et al., 2016) and recent longitudinal imaging analyses suggest a propagation of impairments along the corticofugal tracts (Kassubek et al., 2014; Verstraete et al., 2013). We recently provided the first experimental evidence of the corticofugal hypothesis by showing that absence of SubCerPN, the most important output of the cerebral cortex, delayed disease onset and extended survival in the *Sod1^G86R^* mouse model of the disease (Burg et al., 2019). However, whether corticofugal dissemination of the disease relies on prion-like transmission of misfolded proteins (Braak et al., 2013) or on altered neuronal excitability and subsequent excitotoxicity (Vucic and Kiernan, 2017) remains an open question. Here, we show that ablation of mutant *SOD1^G37R^* transgene from CFuPN had no effect on disease onset and survival. This rules out the possibility of corticofugal propagation of misfolded proteins in a prion-like manner. A few years ago, Thomsen and colleagues knocked-down mutant *SOD1* transgene in the posterior motor cortex of the *SOD1^G93A^* rat model of ALS (Thomsen et al., 2014), using an AAV9 virus that selectively transduces neurons. This strategy delayed disease onset and extended survival. It is possible that the beneficial effect was reached not upon targeting the excitatory CFuPN, but also upon targeting inhibitory interneurons (INs), that modulate excitability and activity of CFuPN and other cortical excitatory neurons (Lodato et al., 2014; Tremblay et al., 2016). In the *TDP-43^A315T^* mouse model of ALS, Zhang and collaborators demonstrated that parvalbumin (PV)-positive INs were hypoactive, while somatostatin (SST)-positive INs were hyperactive {Zhang:2016kw}, and that hyperactivity of SST-positive INs was responsible for hypoactivity of PV-positive INs, and, in turn, for hyperexcitability of layer V SubCerPN. Normal excitability could be restored by genetic ablation of SST-positive INs {Zhang:2016kw}. While the consequences of a knock-down of mutant TDP-43 or *SOD1* on IN excitability is unknown, it is tempting to imagine that the beneficial effect of knocking down *SOD1^G93A^* in the neuronal populations of the motor cortex of transgenic rats arose from the targeting of the INs, in addition to the excitatory neurons, and a possible mitigation of cortical hyperexcitability in these animals. This hypothesis, which remains to be demonstrated, would favour altered neuronal excitability and subsequent excitotoxicity as the detrimental cortical message carried along corticofugal tracts.

### Perspectives

As opposed to the absence of SubCerPN in *Sod1^G86R^* mice (Burg et al., 2019), or the knock-down of *SOD1^G93A^* in the motor cortex of transgenic rats (Thomsen et al., 2014), genetic ablation of *SOD1^G37R^* from CFuPN had no effect on ALS-like symptoms onset and survival. Together, these studies suggest that CFuPN are toxic in a context of ALS, and that their toxicity is not mediated by their cell-autonomous expression of a mutant transgene, but more likely by that of their environment. Because CSN survival and cortical reactive gliosis seemed to be independent from each other, alternative candidates to toxicity could be the populations of INs for their contribution to hyperexcitability (Vucic and Kiernan, 2017). Indeed, cortical hyperexcitability, revealed by transcranial magnetic stimulation, is an early feature of sporadic and familial ALS, appears in *SOD1* mutant carrier prior to clinical onset of the disease (Vucic, 2006), and is negatively correlated with disease progression and survival (Shibuya et al., 2016). Selective silencing of the CFuPN, or selective transgene excision from the INs may in the future contribute to a better understanding of the role of cortical hyperexcitability in ALS, and potentially unravel new therapeutic targets.

## References

1. Arlotta, P., Molyneaux, B.J., Chen, J., Inoue, J., Kominami, R., Macklis, J.D., 2005. Neuronal subtype-specific genes that control corticospinal motor neuron development in vivo. Neuron 45, 207–221. doi:10.1016/j.neuron.2004.12.036

2. Bennett, D.J., Gorassini, M., Sanelli, L., Han, Y., Cheng, J., 1999. Spasticity in rats with sacral spinal cord injury. Journal of Neurotrauma 16, 69–84.

3. Boillee, S., Yamanaka, K., Lobsiger, C.S., Copeland, N.G., Jenkins, N.A., Kassiotis, G., Kollias, G., Cleveland, D.W., 2006. Onset and Progression in Inherited ALS Determined by Motor Neurons and Microglia. Science 312, 1389–1392. doi:10.1126/science.1123511

4. Braak, H., Brettschneider, J., Ludolph, A.C., Lee, V.M., Trojanowski, J.Q., Del Tredici, K., 2013. Amyotrophic lateral sclerosis-a model of corticofugal axonal spread. Nature Reviews Neurology 9, 708–714. doi:10.1038/nrneurol.2013.221

5. Brettschneider, J., Del Tredici, K., Toledo, J.B., Robinson, J.L., Irwin, D.J., Grossman, M., Suh, E., Van Deerlin, V.M., Wood, E.M., Baek, Y., Kwong, L., Lee, E.B., Elman, L., McCluskey, L., Fang, L., Feldengut, S., Ludolph, A.C., Lee, V.M.Y., Braak, H., Trojanowski, J.Q., 2013. Stages of pTDP-43 pathology in amyotrophic lateral sclerosis. Ann Neurol. 74, 20–38. doi:10.1016/S1474-4422(12)70014-5

6. Brown, R.H., Al-Chalabi, A., 2017. Amyotrophic Lateral Sclerosis. N. Engl. J. Med. 377, 162–172. doi:10.1056/NEJMra1603471

7. Burg, T., Bichara, C., Scekic-Zahirovic, J., Fischer, M., Stuart-Lopez, G., Lefebvre, F., Cordero-Erausquin, M., Rouaux, C., 2019. Absence of subcerebral projection neurons delays disease onset and extends survival in a mouse model of ALS. bioRxiv. doi:10.1038/nn.4257

8. Carlson, A.L., Bennett, N.K., Francis, N.L., Halikere, A., Clarke, S., Moore, J.C., Hart, R.P., Paradiso, K., Wernig, M., Kohn, J., Pang, Z.P., Moghe, P.V., 2016. Generation and transplantation of reprogrammed human neurons in the brain using 3D microtopographic scaffolds. Nature Communications 7, 1–10. doi:10.1038/ncomms10862

9. Charcot, J.M., 1874. Sclérose Latérale Amyotrophique: Oeuvres Complètes. Bureaux du Progrès Medical, Paris.

10. Chia, R., Chiò, A., Traynor, B.J., 2017. Novel genes associated with amyotrophic lateral sclerosis: diagnostic and clinical implications. The Lancet Neurology 17, 94–102. doi:10.1016/S1474-4422(17)30401-5

11. Dentel, C., Palamiuc, L., Henriques, A., Lannes, B., Spreux-Varoquaux, O., Gutknecht, L., Rene, F., Echaniz-Laguna, A., Gonzalez de Aguilar, J.-L., Lesch, K.P., Meininger, V., Loeffler, J.-P., Dupuis, L., 2013. Degeneration of serotonergic neurons in amyotrophic lateral sclerosis: a link to spasticity. Brain: a journal of neurology 136, 483–493. doi:10.1016/0379-0738(93)90237-5

12. Eisen, A., Braak, H., Del Tredici, K., Lemon, R., Ludolph, A.C., Kiernan, M.C., 2017. Cortical influences drive amyotrophic lateral sclerosis. Journal of Neurology, Neurosurgery & Psychiatry 88, jnnp-2017–315573. doi:10.1136/jnnp-2017-315573

13. Fame, R.M., MacDonald, J.L., Macklis, J.D., 2011. Development, specification, and diversity of callosal projection neurons. Trends Neurosci. 34, 41–50. doi:10.1016/j.tins.2010.10.002

14. Farrawell, N.E., Lambert-Smith, I.A., Warraich, S.T., Blair, I.P., Saunders, D.N., Hatters, D.M., Yerbury, J.J., 2015. Distinct partitioning of ALSassociated TDP-43, FUS and SOD1mutants into cellular inclusions. Scientific Reports 1–14. doi:10.1038/srep13416

15. Fil, D., DeLoach, A., Yadav, S., Alkam, D., MacNicol, M., Singh, A., Compadre, C.M., Goellner, J.J., O’Brien, C.A., Fahmi, T., Basnakian, A.G., Calingasan, N.Y., Klessner, J.L., Flint Beal, M., Peters, O.M., Metterville, J., Brown, R.H., Jr, Ling, K.K.Y., Rigo, F., Özdinler, P.H., Kiaei, M., 2016. Mutant Profilin1 transgenic mice recapitulate cardinal features of motor neuron disease. Human Molecular Genetics 364, ddw429. doi:10.1523/JNEUROSCI.5253-05.2006

16. Geevasinga, N., Menon, P., Özdinler, P.H., Kiernan, M.C., Vucic, S., 2016. Pathophysiological and diagnosticimplications of cortical dysfunctionin ALS. Nat. Rev. Neurol. 1–11. doi:10.1038/nrneurol.2016.140

17. Guo, W., Fumagalli, L., Prior, R., Van Den Bosch, L., 2017. Current Advances and Limitations in Modeling ALS/FTD in a Dish Using Induced Pluripotent Stem Cells. Frontiers in Neuroscience 11, 1282. doi:10.1073/pnas.1315438110

18. Hardiman, O., Al-Chalabi, A., Brayne, C., Beghi, E., van den Berg, L.H., Chiò, A., Martin, S., Logroscino, G., Rooney, J., 2017. The changing picture of amyotrophic lateral sclerosis: lessons from European registers. Journal of Neurology, Neurosurgery & Psychiatry 88, 557– 563. doi:10.1007/s00415-003-1026-z

19. Herdewyn, S., Cirillo, C., Van Den Bosch, L., Robberecht, W., Berghe, P.V., Van Damme, P., 2014. Prevention of intestinal obstruction reveals progressive neurodegeneration in mutant TDP-43(A315T) mice 9, 1–14. doi:10.1186/1750-1326-9-24

20. Ilieva, H., Polymenidou, M., Cleveland, D.W., 2009. Non-cell autonomous toxicity in neurodegenerative disorders: ALS and beyond. The Journal of Cell Biology 187, 761–772. doi:10.1083/jcb.200908164

21. Ivanhoe, C.B., Reistetter, T.A., 2004. Spasticity. American Journal of Physical Medicine & Rehabilitation 83, S3–S9. doi:10.1097/01.PHM.0000141125.28611.3E

22. Kassubek, J., Muller, H.P., Del Tredici, K., Brettschneider, J., Pinkhardt, E.H., Lule, D., Bohm, S., Braak, H., Ludolph, A.C., 2014. Diffusion tensor imaging analysis of sequential spreading of disease in amyotrophic lateral sclerosis confirms patterns of TDP-43 pathology. Brain: a journal of neurology 137, 1733–1740. doi:10.1517/17530059.2010.536836

23. Lee, J., Hyeon, S.J., Im, H., Ryu, H., Kim, Y., Ryu, H., 2016. Astrocytes and Microglia as Non-cell Autonomous Players in the Pathogenesis of ALS. Exp Neurobiol 25, 233. doi:10.2174/0929867321666140706131825

24. Liu, Y., Pattamatta, A., Zu, T., Reid, T., Bardhi, O., Borchelt, D.R., Yachnis, A.T., Ranum, L.P.W., 2016. C9orf72 BAC Mouse Model with Motor Deficits and Neurodegenerative Features of ALS/FTD. Neuron 90, 521–534. doi:10.1016/j.neuron.2016.04.005

25. Lodato, S., Rouaux, C., Quast, K.B., Jantrachotechatchawan, C., Studer, M., Hensch, T.K., Arlotta, P., 2011. Excitatory projection neuron subtypes control the distribution of local inhibitory interneurons in the cerebral cortex. Neuron 69, 763–779. doi:10.1016/j.neuron.2011.01.015

26. Lodato, S., Shetty, A.S., Arlotta, P., 2014. Cerebral cortex assembly: generating and reprogramming projection neuron diversity. Trends Neurosci. 38, 117–125. doi:10.1016/j.tins.2014.11.003

27. Lutz, C., 2018. Brain Research. Brain Research 1693, 1–10. doi:10.1016/j.brainres.2018.03.024

28. Madisen, L., Zwingman, T.A., Sunkin, S.M., Oh, S.W., Zariwala, H.A., Gu, H., Ng, L.L., Palmiter, R.D., Hawrylycz, M.J., Jones, A.R., Lein, E.S., Zeng, H., 2009. A robust and high-throughput Cre reporting and characterization system for the whole mouse brain. Nat. Neurosci. 13, 133–140. doi:10.1038/nprot.2006.58

29. Marques, C., Fischer, M., Keime, C., Burg, T., Brunet, A., Scekic-Zahirovic, J., Rouaux, C., 2019. Early alterations of RNA metabolism and splicing from adult corticospinal neurons in an ALS mouse model. bioRxiv. doi:10.1101/gr.1239303

30. Molyneaux, B.J., Arlotta, P., Hirata, T., Hibi, M., Macklis, J.D., 2005. Fezl is required for the birth and specification of corticospinal motor neurons. Neuron 47, 817–831. doi:10.1016/j.neuron.2005.08.030

31. Molyneaux, B.J., Arlotta, P., Menezes, J.R.L., Macklis, J.D., 2007. Neuronal subtype specification in the cerebral cortex. Nat. Rev. Neurosci. 8, 427–437. doi:10.1038/nrn2151

32. Müller, H.-P., Brenner, D., Roselli, F., Wiesner, D., Abaei, A., Gorges, M., Danzer, K.M., Ludolph, A.C., Tsao, W., Wong, P.C., Rasche, V., Weishaupt, J.H., Kassubek, J., 2019. Longitudinal diffusion tensor magnetic resonance imaging analysis at the cohort level reveals disturbed cortical and callosal microstructure with spared corticospinal tract in the TDP-43 1–13. doi:10.1186/s40035-019-0163-y

33. Nihei, K., Kowall, N.W., 1993. Involvement of NPY-immunoreactive neurons in the cerebral cortex of amyotrophic lateral sclerosis patients. Neuroscience Letters 1–4.

34. Oussini, El, H., Scekic-Zahirovic, J., Vercruysse, P., Marques, C., Dirrig-Grosch, S., Dieterlé, S., Picchiarelli, G., Sinniger, J., Rouaux, C., Dupuis, L., 2017. Degeneration of serotonin neurons triggers spasticity in amyotrophic lateral sclerosis. Ann Neurol. 82, 444–456. doi:10.1038/nn.3544

35. Ozdinler, P.H., Benn, S., Yamamoto, T.H., Guzel, M., Brown, R.H., Macklis, J.D., 2011. Corticospinal Motor Neurons and Related Subcerebral Projection Neurons Undergo Early and Specific Neurodegeneration in hSOD1G93A Transgenic ALS Mice. The Journal of Neuroscience 31, 4166–4177. doi:10.1523/JNEUROSCI.4184-10.2011

36. Purves, D., Augustine, G.J., Fitzpatrick, D., Hall, W.C., LaMantia, A.-S., McNamara, J.O., Williams, S.M., 2004. Neuroscience. Sinauer Associates, Inc. 1–832.

37. Ripps, M.E., Huntley, G.W., Hof, P.R., Morrison, J.H., Gordon, J.W., 1995. Transgenic mice expressing an altered murine superoxide dismutase gene provide an animal model of amyotrophic lateral sclerosis. Proc. Natl. Acad. Sci. U.S.A. 92, 689–693.

38. Rouaux, C., Panteleeva, I., Rene, F., Gonzalez de Aguilar, J.-L., Echaniz-Laguna, A., Dupuis, L., Menger, Y., Boutillier, A.-L., Loeffler, J.-P., 2007. Sodium valproate exerts neuroprotective effects in vivo through CREB-binding protein-dependent mechanisms but does not improve survival in an amyotrophic lateral sclerosis mouse model. The Journal of Neuroscience: the official journal of the Society for Neuroscience 27, 5535–5545. doi:10.1523/JNEUROSCI.1139-07.2007

39. Scekic-Zahirovic, J., Oussini, El, H., Mersmann, S., Drenner, K., Wagner, M., Sun, Y., Allmeroth, K., Dieterlé, S., Sinniger, J., Dirrig-Grosch, S., Rene, F., Dormann, D., Haass, C., Ludolph, A.C., Lagier-Tourenne, C., Storkebaum, E., Dupuis, L., 2017. Motor neuron intrinsic and extrinsic mechanisms contribute to the pathogenesis of FUS-associated amyotrophic lateral sclerosis. Acta Neuropathol. 1–20. doi:10.1007/s00401-017-1687-9

40. Serio, A., Patani, R., 2017. Concise Review: The Cellular Conspiracy of Amyotrophic Lateral Sclerosis. Stem Cells 36, 293–303. doi:10.1038/nm.4052

41. Shibuya, K., Park, S.B., Geevasinga, N., Menon, P., Howells, J., Simon, N.G., Huynh, W., Noto, Y.-I., Götz, J., Kril, J.J., Ittner, L.M., Hodges, J., Halliday, G., Vucic, S., Kiernan, M.C., 2016. Motor cortical function determines prognosis in sporadic ALS. Neurology 87, 513–520. doi:10.1212/WNL.0000000000002912

42. Sorenson, E.J., 2012. The Electrophysiology of the Motor Neuron Diseases. Neurologic Clinics 30, 605–620. doi:10.1016/j.ncl.2011.12.006

43. Thomsen, G.M., Gowing, G., Latter, J., Chen, M., Vit, J.P., Staggenborg, K., Avalos, P., Alkaslasi, M., Ferraiuolo, L., Likhite, S., Kaspar, B.K., Svendsen, C.N., 2014. Delayed Disease Onset and Extended Survival in the SOD1G93A Rat Model of Amyotrophic Lateral Sclerosis after Suppression of Mutant SOD1 in the Motor Cortex. Journal of Neuroscience 34, 15587– 15600. doi:10.1523/JNEUROSCI.2037-14.2014

44. Tremblay, R., Lee, S., Rudy, B., 2016. GABAergic Interneurons in the Neocortex: From Cellular Properties to Circuits. Neuron 91, 260–292. doi:10.1016/j.neuron.2016.06.033

45. van Es, M.A., Hardiman, O., Chiò, A., Al-Chalabi, A., Pasterkamp, R.J., Veldink, J.H., van den Berg, L.H., 2017. Amyotrophic lateral sclerosis. The Lancet 390, 2084–2098. doi:10.1016/S0140-6736(17)31287-4

46. Vergouts, M., Marinangeli, C., Ingelbrecht, C., Genard, G., Schakman, O., Sternotte, A., Calas, A.-G., Hermans, E., 2015. Early ALS-type gait abnormalities in AMP-dependent protein kinase-deficient mice suggest a role for this metabolic sensor in early stages of the disease. Metab Brain Dis 30, 1369–1377. doi:10.1177/0300060514554725

47. Verstraete, E., Veldink, J.H., van den Berg, L.H., van den Heuvel, M.P., 2013. Structural brain network imaging shows expanding disconnection of the motor system in amyotrophic lateral sclerosis. Hum. Brain Mapp. 35, 1351–1361. doi:10.1016/j.neuron.2012.03.004

48. Vucic, S., 2006. Novel threshold tracking techniques suggest that cortical hyperexcitability is an early feature of motor neuron disease. Brain: a journal of neurology 129, 2436–2446. doi:10.1093/brain/awl172

49. Vucic, S., Kiernan, M.C., 2017. Transcranial Magnetic Stimulationfor the Assessment of Neurodegenerative Disease. Neurotherapeutics 1–16. doi:10.1007/s13311-016-0487-6

50. Wegorzewska, I., Bell, S., Cairns, N.J., Miller, T.M., Baloh, R.H., 2009. TDP-43 mutant transgenic mice develop features of ALS and frontotemporal lobar degeneration. PNAS 106, 18809– 18814.

51. Weskamp, K., Tank, E.M., Miguez, R., McBride, J.P., Gómez, N.B., White, M., Lin, Z., Moreno Gonzalez, C., Serio, A., Sreedharan, J., Barmada, S.J., 2019. Shortened TDP43 isoforms upregulated by neuronal hyperactivity drive TDP43 pathology in ALS. J. Clin. Invest. doi:10.1172/JCI130988DS1

52. Yasvoina, M.V., Genc, B., Jara, J.H., Sheets, P.L., Quinlan, K.A., Milosevic, A., Shepherd, G.M.G., Heckman, C.J., Ozdinler, P.H., 2013. eGFP Expression under UCHL1 Promoter Genetically Labels Corticospinal Motor Neurons and a Subpopulation of Degeneration-Resistant Spinal Motor Neurons in an ALS Mouse Model. Journal of Neuroscience 33, 7890–7904. doi:10.1523/JNEUROSCI.2787-12.2013

53. Zang, D.W., Cheema, S.S., 2002. Degeneration of corticospinal and bulbospinal systems in the superoxide dismutase 1. Neuroscience Letters 332, 99–102.

